# Bright near-infrared genetically encoded voltage indicator for all-optical electrophysiology

**DOI:** 10.1101/536359

**Authors:** Mikhail V. Monakhov, Mikhail E. Matlashov, Michelangelo Colavita, Chenchen Song, Daria M. Shcherbakova, Srdjan D. Antic, Vladislav V. Verkhusha, Thomas Knöpfel

## Abstract

We developed genetically encoded voltage indicators (GEVIs) using bright near-infrared (NIR) fluorescent proteins from bacterial phytochromes. These new NIR GEVIs are optimized for combination of voltage imaging with simultaneous blue light optogenetic actuator activation. Iterative optimizations led to a GEVI here termed nirButterfly, which reliably reports neuronal activities including subthreshold membrane potential depolarization and hyperpolarization, as well as spontaneous spiking, or electrically- and optogenetically-evoked action potentials. This enables largely improved all-optical causal interrogations of physiology.

---

Development of increasingly better performing genetically encoded voltage indicators (GEVIs) has been ongoing for the last several decades^1–3^. Current GEVIs already allow the direct optical monitoring of fast electrical signals at high spatiotemporal resolution and coverage *in vitro* and *in vivo*^4–7^. Like genetically encoded calcium indicators (GECIs), GEVIs enable cell class specific large-scale mapping of neuronal activities *in vivo*^1, 8–10^. In contrast to GECIs, GEVIs are not only able to report action potentials, but also depolarizing subthreshold potentials and hyperpolarizing (inhibitory) potentials that underlie information processing at the neuronal level. GEVIs also supersede classic voltage-sensitive dyes, which are difficult to target to specific cell classes and require invasive staining procedures. GEVI-based voltage imaging therefore offers enormous potential for delineating functional electrical processing mechanisms in neurophysiology.

Both observation and control of electrical signals of defined neuronal populations is necessary to establish causal relationships between neuronal circuit mechanism and behaviors. This can be achieved simultaneously by the combined expression of spectrally non-overlapping optogenetic actuators for stimulating activity and GEVIs for recording the activity; an approach that has been coined as “all-optical electrophysiology” (AOE)^11^. Current AOE offers proof-of-principle but is limited by the low fluorescence quantum yield (QY) of the optical indicators of the QuasAr/Archon type^11, 12^. High excitation intensities required to generate sufficient fluorescence output from these opsins restrict their application due to requirement of very intense light sources and concerns for tissue heating^13^.

To overcome these technical limitations, we used monomeric near-infrared (NIR) fluorescent proteins of the miRFP family engineered from bacterial phytochromes, which offer the same spectral range as the opsin-based GEVIs QuasAr^11^ and Archon^12^, but with much higher QY (i.e. they are ‘brighter’; **Supplementary Table**)^14^. We anticipated that the segregated spectral ranges of miRFPs and the cation channel opsin CheRiff^11^ (**Fig. 1a**) allow largely improved AOE with high application potential in biological systems.

**Figure 1.**
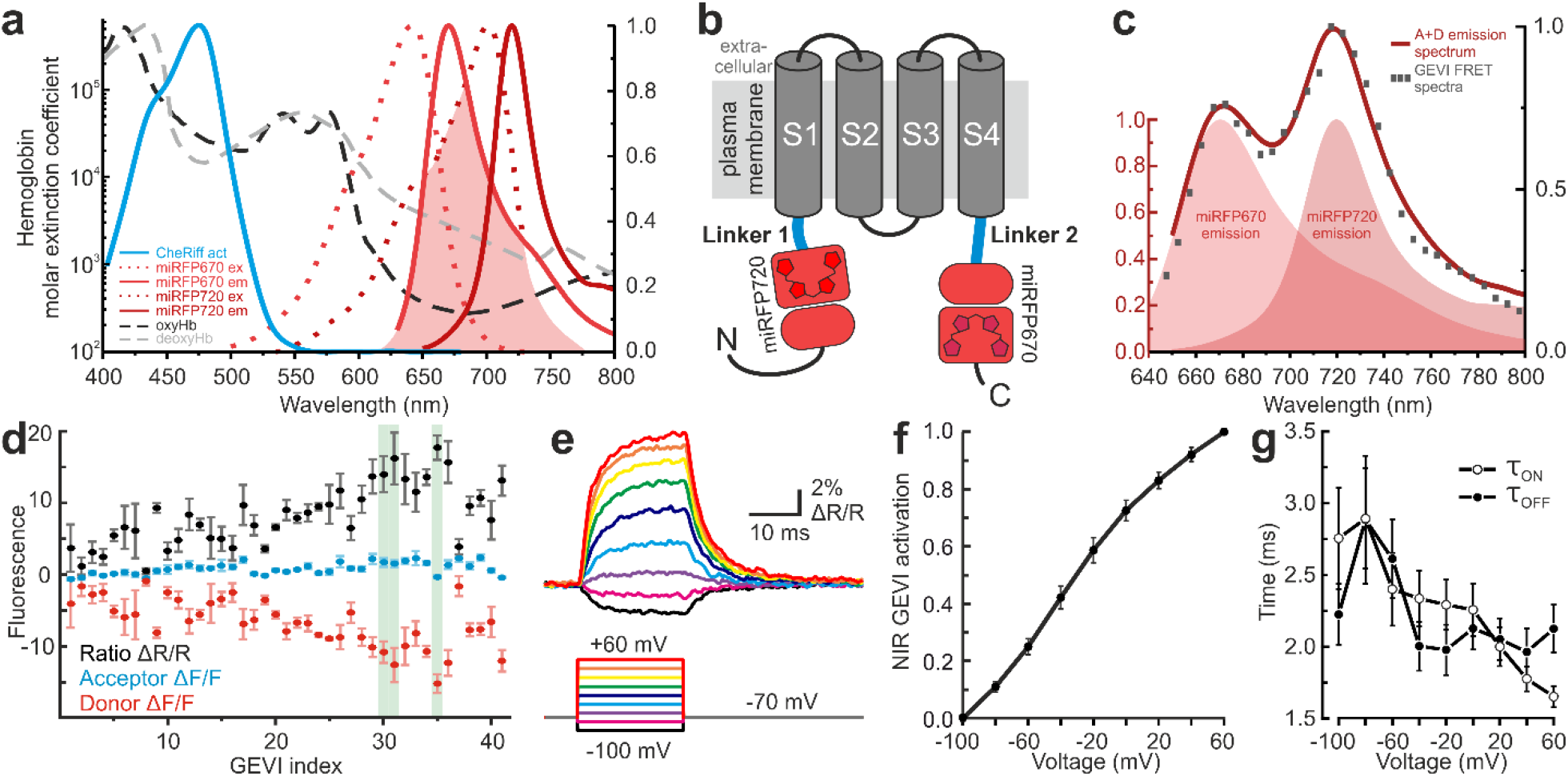
NIR GEVI design and functional characterization. (**a**) Non-overlapping spectral compatibility between the excitation (ex) and emission (em) spectra of miRFPs (500-800 nm) with the activation spectrum (act) of the blue-shifted cation channel opsin CheRiff. Shaded area indicates the overlap integral between miRFP670 (FRET donor) emission and miRFP720 (FRET acceptor) excitation to enable FRET. Spectral bands of miRFPs lie in a region with low extinction coefficient (cm^−1^ M^−1^) of oxyhemoglobin (oxyHb) and deoxyhemoglobin (deoxyHb). (**b**) Schematic of the NIR GEVI structural design. A synthetic voltage sensing domain derived from the *Ciona intestinalis* voltage sensing phosphatase and Kv3.1 potassium channel is sandwiched between a FRET pair of miRFPs – miRFP670 (donor) and miRFP720 (acceptor). The two linker regions highlighted in blue were varied during evolution towards better performing variants. (**c**) Emission spectra from nirButterfly (excitation at 633 nm) (black squares) in comparison to the unweighted sum of donor and acceptor emission spectra. (**d-g**) Functional characterization of NIR GEVIs in voltage clamped HEK293 cells. (**d**) Evolution of NIR GEVIs. Fluorescence signals against index of NIR GEVI variants (index) interactively generated and tested. Green shading highlights selected best-performing first generation NIR GEVI variants. (**e**) Representative recording of fluorescence emission ratio from nirButterfly for a family of voltage steps (20 ms, from −100 mV to +60 mV in 20-mV step size) from a −70 mV holding potential. Average over 10 trials. (**f**) nirButterfly normalized voltage-activation dependency (N = 15 cells; mean ± SD). (**g**) nirButterfly ON- and OFF-response time constants, from single exponential fit (N = 9 cells for τ_ON_, 8 cells for τ_OFF_; mean ± SEM; **Supplementary Figure 2**).

We generated ratiometric NIR GEVIs from voltage sensitive fluorescent protein (VSFP) chimeric Butterfly design - using a synthetic voltage sensing domain derived from the *Ciona intestinalis* voltage sensing phosphatase^15^ and Kv3.1 potassium channel (Ci-VSD)^16^ - by substituting its Forster resonance energy transfer (FRET) fluorescent protein (FP) pair with a pair of miRFPs. (miRFP670^14^ and miRFP720, FRET donor and FRET acceptor, respectively; **Fig. 1b**). Early prototypes of these NIR GEVIs exhibited FRET (**Fig. 1c**) and retained the localized membrane targeting properties from VSFPs when expressed in cultured human embryonic kidney-293 (HEK293) cells (**Supplementary Fig. 1a-b**). Functional testing demonstrated a decrease in donor fluorescence and a smaller increase of acceptor fluorescence upon membrane depolarization of patch-clamped HEK293 cells (**Fig. 1d, Supplementary Fig. 1b-d**). This feature allows ratiometric measurements (**acceptor/donor**) known to be more resistant to artefacts from movement and dye fading compared with monochromatic indicators. We successively evolved better performing variants (larger dynamic range with fast kinetics) by iterating between patch clamp assay and variations in both the length and amino acid composition of the linker regions between donor FP and VSD and between VSD and acceptor FP (Linkers 1 and 2 respectively, **Figs. 1b, 1d, Supplementary Fig. 1**). We then selected the three best candidate variants (dubbed NIR GEVIa, −b –c, highlighted green in **Fig. 1d**) for a more thorough biophysical characterization in HEK293 cells (**Figs. 1e-g, Supplementary Fig. 2**). In this assay, the fluorescence ratio of NIR GEVIb increased by up to about 8% (5.58% ± 2.16 ΔR/R; mean ± SD, N = 11 cells) upon a depolarization step from −80 mV to 20 mV, with an approximately linear relationship between fluorescence ratio and membrane voltage (**Fig. 1f**). Importantly, the response ON- and OFF-time constants were fast (2.16 ± 0.55 ms, from −70 mV to 0 mV; 2.13 ± 0.42 ms, from 0 mV to −70 mV when fitted with a single exponential time constant; mean ± SD) and time constants were less voltage-dependent than those observed with yellow/red Butterfly^16^ (**Fig. 1g, Supplementary Fig. 2**).

NIR GEVIa, −b –c were then evaluated in neurons of cortical primary cultures. All three exhibited very good sensitivity (NIR GEVIa: 9.3 ± 1.2 % ΔR/R / 100 mV, NIR GEVIb: 9.3 ± 2.6 % ΔR/R / 100 mV, NIR GEVIc 5.6 ± 1.1% ΔR/R / 100 mV). We have chosen NIR GEVIb as the most promising variant and termed it nirButterfly. nirButterfly reports action potentials and subthreshold membrane potential fluctuations with single sweep resolution (**Figs. 2b-c**) using relatively modest excitation intensity, not exceeding 50 mW/mm^2^, a value much below the average power density of infrared light routinely used for two photon microscopy.

**Figure 2.**
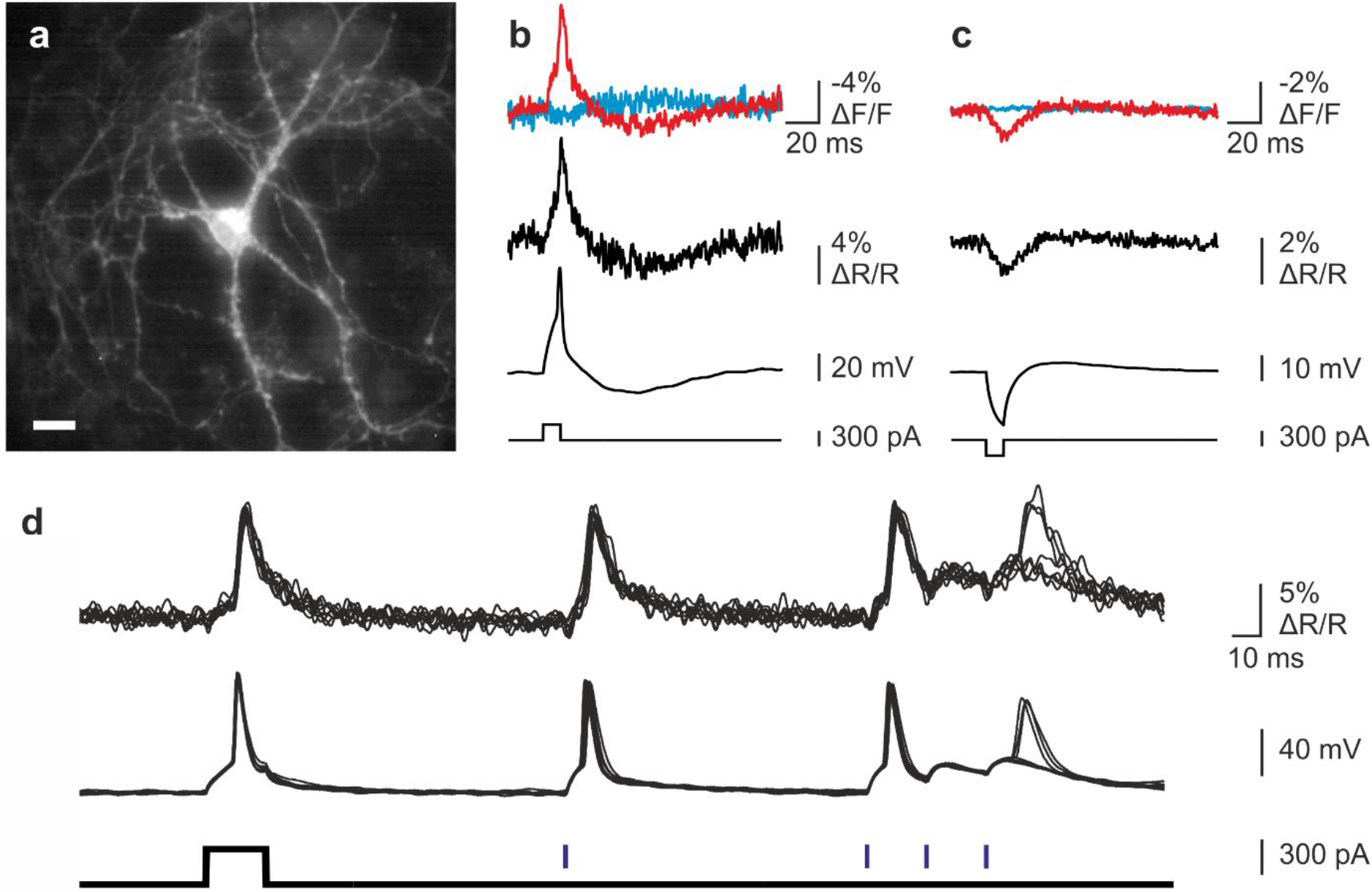
All-optical electrophysiology in neurons. (**a**) Epifluorescence image of a cultured neuron expressing NIR GEVIa. Scale bar = 10 μm (**b**) Optical monitoring of evoked (300 pA current injection) action potential from the current-clamped neuron shown in a. FRET donor (red) and acceptor (blue) signals provide a ratiometric (middle, black) report of the microelectrode measurements (lower, black) with high fidelity (average of 7 trials shown). (**c**) NIR GEVI resolves current injection-induced hyperpolarization (average of 20 trials shown). Traces as in **b**. (**d**) Neuron co-expressing nirButterfly and the cation channel opsin CheRiff shows nirButterfly-enabled all-optical electrophysiology. nirButterfly resolves microelectrode current injection-evoked and CheRiff photocurrent-triggered action potentials and subthreshold depolarizations. Upper traces show optical signals (8 individual trials superimposed), middle traces show the microelectrode recordings, lower traces show microelectrode (black) and blue light (purple) pulses.

Finally, we established AOE by co-transfecting cultured cortical neurons with a nirButterfly plasmid and a CheRiff-IRES-NLS-mTagBFP2 plasmid encoding the blue-shifted cation channel opsin variant CheRiff. Like current injections through a patch clamp electrode, the CheRiff-mediated photocurrent triggered action potentials and subthreshold depolarizations were readily resolved in the optical recording (**Figs. 2d**). Most importantly, the red light used for nirButterfly fluorescence excitation did not induce any changes in membrane potential, as expected from the complete spectral separation of the actuator opsin and NIR GEVI (**Fig. 1a**). NIR GEVIs offer sufficient SNR for reliable detection of action potential without averaging (**Fig. 2d, Supplementary Fig. 3**).

nirButterfly outperforms currently available red and NIR GEVIs (for comparison see **Supplementary Table**) and enables improved AOE, with enhanced quantum yield that allows use of modest red light excitation intensity suitable for biological applications. Imaging in the NIR spectral range brings additional benefits: 1) A bypass of tissue auto-fluorescence and hemoglobin signals; 2) An increased tissue penetration for deep-tissue imaging^17^; and 3) Two-photon voltage imaging in the NIR range using Soret band activation of the chromophore biliverdin^18^. AOE using nirButterfly will improve the experimental studies of causal physiological mechanisms in the living systems.

## Acknowledgments

We thank all members of the Knöpfel, Verkhusha and Antic labs for support and discussions. We thank Adam Cohen for CheRiff spectra. This work is supported by NIH Brain Initiative grants U01 NS099573, U01-MH019091 and R01 GM122567.

## Author contributions

All authors collaborated on protein design. MEM, MVM, DMS, CS and MC performed molecular cloning, GEVI evolution, biophysical characterization and neurophysiological experiments. TK, VVV and SDA directed the research. CS and TK wrote manuscript and prepared figures with input from all authors.

## Availability of data

Data are available from the corresponding authors upon reasonable request. **Accession codes** NIR GEVI nucleotide sequence: Genbank XXXXXNIR GEVI expression plasmid: Addgene XXXXX

## METHODS

### Plasmids

Fluorescent proteins used in chimeric VSFP Butterfly^16^ were substituted by NIR FPs miRFP670 and miRFP720^14^ at the C- and N-termini respectively of the synthetic voltage sensing domain derived from *Ciona intestinalis* voltage sensing phosphatase and Kv3.1 potassium channel (Ci-VSD^19^; **Fig. 1b**). Using this scaffold, a set of constructs with modifications of Linker 1 (between miRFP720 and Ci-VSD) or Linker 2 (between Kv3.1 and miRFP670) were generated either by overlap extension PCR or using site-directed mutagenesis (QuickChange, Agilent, USA). Plasmid pPuro-CAG-CheRiff-IRES-NLS-mTagBFP2 encoding the optogenetic activating opsin CheRiff was made by subcloning CheRiff, IRES and NLS-mTagBFP2 into pcDNA3.1-Puro-CAG-ASAP vector.

### NIR GEVI screening in HEK cells

HEK293 cells were maintained in DMEM supplemented with 10% FBS, 2 mM glutamine, 100 U/ml penicillin and 100 μg/ml streptomycin. Cells were transfected using Lipofectamine 2000 in 24-well plates, and were seeded onto poly-D-lysine coated coverslips two days after transfection. GEVI testing was performed one day after seeding to coverslips. Biliverdin was added one day before recording to a final concentration of 25 μM. Cells were washed with DPBS and placed in recording chamber with external solution containing (in mM): 125 NaCl, 0.5 KCl, 1 MgCl_2_, 3 CaCl_2_, 30 glucose, 1 octanol and 10 HEPES, pH 7.3, 305 mOsm. Internal solution contained (in mM): 125 K gluconate, 8 NaCl, 0.6 MgCl_2_, 0.1 CaCl_2_, 1 EGTA, 4 MgATP, 0.4 NaGTP and 10 HEPES, pH 7.3, 295 mOsm. Osmolarity was adjusted with sucrose. Glass pipettes (1.5/0.75 OD/ID) were pulled with Flaming/Brown micropipette puller P-97 to a resistance of 6-10 MΩ. Recordings were performed at room temperature. In HEK cells the voltage was changed (from the holding potential of −70 mV) to −100, −30, 30 and 100 mV in voltage clamp mode using MultiClamp700B amplifier and Digidata1440A digitizer (Molecular Devices, San Jose, CA). Duration of each step and each period between steps was 500 ms. An LED light source with GYR (‘Green-Yellow-Red’) module was used for excitation of fluorophores (CoolLED, UK). Images were acquired using Olympus BX51WI microscope, NeuroCCD camera (gain 30x, 80×80 pixels, RedShirtImaging, Decatur, GA), 605/30 nm excitation filter (ET605/30x), 640 nm dichroic mirror (T640lpxr), 667/40 nm (ET667/30m) and 720/40 nm (ET720/40m) emission filters. ΔF/F values were calculated for each variant after imaging at frame rate of 20 Hz and kinetics parameters were estimated after imaging at 500 Hz.

### Biophysical characterization of NIR GEVI variants in HEK cells

HEK293 cells were transfected with NIR GEVI variants using lipofectamine, with biliverdin added at time of transfection. Cells were re-plated on poly-D-lysine coated coverslips at least 8 hours after transfection. Whole cell recordings were performed as previously described^4,5^ at 32-34°C, where a family of voltage steps (20 ms; from −100 mV to +60 mV) from −70 mV holding potential was applied to an indicator-expressing voltage-clamped HEK293 cell. Fluorescence was recorded using photodiodes (TILL Photonics, Gräfelfing, Germany) at 10 kHz. The acceptor/donor signal was fitted with a single exponential function for estimation of the ON- and OFF-response time constants.

GEVI FRET spectra was measured using Leica TCS SP5 equipped with 40X (NA 0.85) objective using 633 nm He-Ne laser for NIR GEVI excitation, fluorescence was imaged in the range of 640 to 800 nm, at 10 nm bandwidth in 5 nm λ steps. ROIs were manually defined over the expressing HEK cells.

### Animals

All procedures were in accordance with the UK Animal Scientific Procedures Act (1986) under Home Office Project and Personal Licences following appropriate ethical review.

### Neuronal all-optical electrophysiology

Primary cultures of cortical and hippocampal neurons were dissociated from P0.5 C57BL/6 mice and plated onto coverslips pre-coated with poly-D-lysine, and maintained in Neurobasal medium supplemented with 2 mM L-glutamine, 0.5% B-27 and 1% penicillin/streptomycin^20^. Cells were transfected 4 days after plating using calcium phosphate^21^ (Invitrogen), exogenous biliverdin (Sigma) was added 2 days after transfection.

Experiments were performed 14-16 days after transfection. Coverslips with transfected cells were placed in a recording chamber mounted on the stage of a dual emission widefield epifluorescence immersion microscope (Scientifica, UK) and superfused with external solution with the following composition (in mM): 110 NaCl, 3 KCl, 1 MgSO_4_, 3 CaCl_2_, 10 glucose, 0.5 Na_2_HPO_4_ and 10 HEPES, adjusted to pH 7.4 with NaOH (23°C). Whole-cell current clamp recordings were performed with borosilicate glass pipettes (5-7 MΩ) pulled on a two-stage vertical puller (PC-10, Narishige, Japan), and filled with internal solution containing (in mM): 120 K gluconate, 20 KCl, 3 MgCl_2_, 0.8 CaCl_2_, 2 EGTA, 4.4 MgATP and 15 HEPES, adjusted to pH 7.3 with KOH. Signals were acquired with an Axon 700B Multiclamp amplifier and digitized at 10 kHz with a Digidata 1440A using pCLAMP software (Molecular Devices, San Jose, CA).

The microscope is equipped with two synchronised CMOS cameras (acA1920-155um, Basler AG), or photodiodes (TILL Photonics, Gräfelfing, Germany), using LED light sources (CoolLED, UK; FiberOptoMeter, NPI Electronic, Germany) and the following optical filters (Semrock and Chroma) for NIR GEVI voltage imaging: miRFP670 (donor) excitation FF02-615/20, miRFP670 emission 680/60, miRFP720 emission FF01-775/140, excitation beamsplitter 635LP (FF635-Di01), and detection beamsplitter 700LP (FF700-Di01). CheRiff was activated by blue band pass filtered (FF01-482/35) LED light which was combined with the red LED excitation light using a 600 nm long pass dichroic mirror.

For expression characterization, cells were fixed using 2% paraformaldehyde after optical imaging experiments, and imaged using Leica TCS SP5 equipped with 63X (NA 1.40) oil immersion objective using 633 nm He-Ne laser for miRFP (NIR GEVI) excitation, and 405nm for NLS-mTagBFP2 (CheRiff) excitation.

### Data analysis

Optical and electrophysiological signals were analyzed using Clampfit (Molecular Devices) and custom programs under Matlab 7.4 (MathWorks) and Origin 8 software (OriginLab). Ratiometric calculations were performed as previously described^4, 5^. Analysis codes are available upon reasonable request.

## References

1. Knopfel, T. Genetically encoded optical indicators for the analysis of neuronal circuits. Nat Rev Neurosci 13, 687–700 (2012).

2. Lin, M.Z. & Schnitzer, M.J. Genetically encoded indicators of neuronal activity. Nat Neurosci 19, 1142–1153 (2016).

3. Antic, S.D., Empson, R.M. & Knopfel, T. Voltage imaging to understand connections and functions of neuronal circuits. J Neurophysiol 116, 135–152 (2016).

4. Akemann, W. et al. Imaging neural circuit dynamics with a voltage-sensitive fluorescent protein. J Neurophysiol 108, 2323–2337 (2012).

5. Akemann, W., Mutoh, H., Perron, A., Rossier, J. & Knopfel, T. Imaging brain electric signals with genetically targeted voltage-sensitive fluorescent proteins. Nat Methods 7, 643–649 (2010).

6. Yang, H.H. et al. Subcellular Imaging of Voltage and Calcium Signals Reveals Neural Processing In Vivo. Cell 166, 245–257 (2016).

7. Gong, Y. et al. High-speed recording of neural spikes in awake mice and flies with a fluorescent voltage sensor. Science 350, 1361–1366 (2015).

8. Carandini, M. et al. Imaging the awake visual cortex with a genetically encoded voltage indicator. J Neurosci 35, 53–63 (2015).

9. Allen, W.E. et al. Global Representations of Goal-Directed Behavior in Distinct Cell Types of Mouse Neocortex. Neuron 94, 891–907 e896 (2017).

10. Madisen, L. et al. Transgenic mice for intersectional targeting of neural sensors and effectors with high specificity and performance. Neuron 85, 942–958 (2015).

11. Hochbaum, D.R. et al. All-optical electrophysiology in mammalian neurons using engineered microbial rhodopsins. Nat Methods 11, 825–833 (2014).

12. Piatkevich, K.D. et al. A robotic multidimensional directed evolution approach applied to fluorescent voltage reporters. Nat Chem Biol (2018).

13. Lou, S. et al. Genetically Targeted All-Optical Electrophysiology with a Transgenic Cre-Dependent Optopatch Mouse. J Neurosci 36, 11059–11073 (2016).

14. Shcherbakova, D.M. et al. Bright monomeric near-infrared fluorescent proteins as tags and biosensors for multiscale imaging. Nat Commun 7, 12405 (2016).

15. Murata, Y., Iwasaki, H., Sasaki, M., Inaba, K. & Okamura, Y. Phosphoinositide phosphatase activity coupled to an intrinsic voltage sensor. Nature 435, 1239–1243 (2005).

16. Mishina, Y., Mutoh, H., Song, C. & Knopfel, T. Exploration of genetically encoded voltage indicators based on a chimeric voltage sensing domain. Front Mol Neurosci 7, 78 (2014).

17. Xu, Y., Zou, P. & Cohen, A.E. Voltage imaging with genetically encoded indicators. Curr Opin Chem Biol 39, 1–10 (2017).

18. Piatkevich, K.D. et al. Near-Infrared Fluorescent Proteins Engineered from Bacterial Phytochromes in Neuroimaging. Biophys J 113, 2299–2309 (2017).

19. Mishina, Y., Mutoh, H. & Knopfel, T. Transfer of Kv3.1 voltage sensor features to the isolated Ci-VSP voltage-sensing domain. Biophys J 103, 669–676 (2012).

20. Beaudoin, G.M., 3rd et al. Culturing pyramidal neurons from the early postnatal mouse hippocampus and cortex. Nat Protoc 7, 1741–1754 (2012).

21. Jiang, M. & Chen, G. High Ca2+-phosphate transfection efficiency in low-density neuronal cultures. Nat Protoc 1, 695–700 (2006).

