## Supplemental Information for "Bright near-infrared genetically encoded voltage indicator for all-optical electrophysiology"

### SUPPLEMENTARY INFORMATION

**Overview**

| | |
|---|---|
| **Supplementary Fig. 1** | NIR GEVI variant screening in HEK293 cells |
| **Supplementary Fig. 2** | NIR GEVI biophysical characterisation in HEK293 cells |
| **Supplementary Fig. 3** | AOE demonstrated in neuron co-expressing NIR GEVIa and CheRiff |
| **Supplementary Table** | Main characteristics of current red-shifted GEVIs |

### Supplementary figures

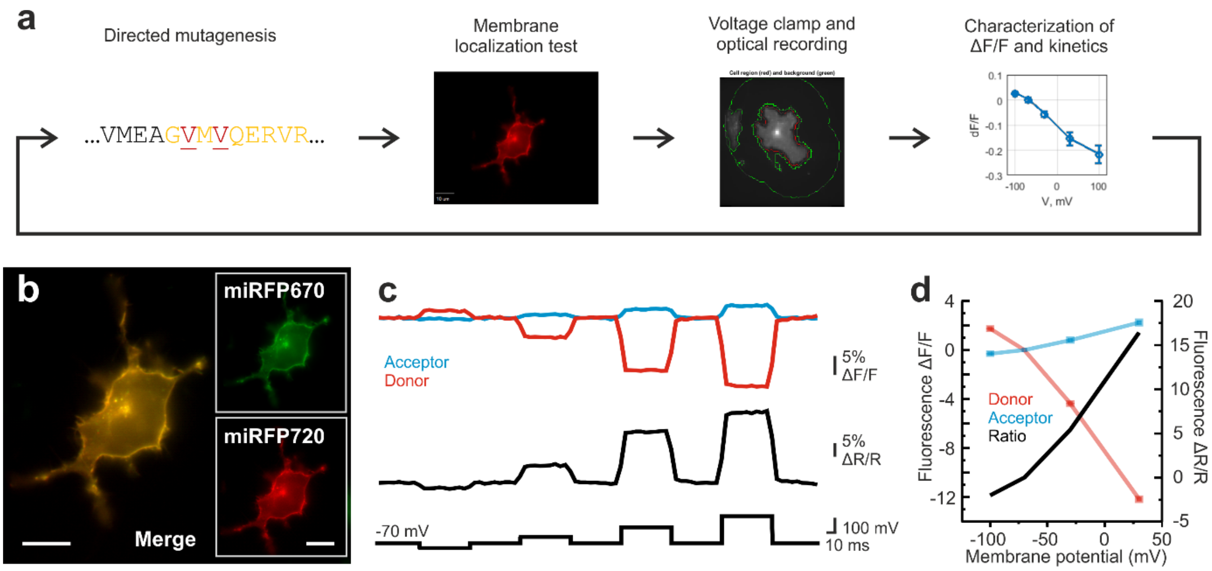

**Supplementary Figure 1.** NIR GEVI variant screening in HEK293 cells. (**a**) The workflow for screening of GEVI variants in HEK293 cells. (**b**) NIR GEVI variants retained the localised membrane targeting properties from VSFPs when expressed in cultured HEK293 cells, through both donor (miRFP670) and acceptor (miRFP720) FP channels. Scale bar = 10μm. (**c**) In the NIR GEVI screening protocol, selected voltage steps from -70 mV holding potential are applied to the patch-clamped cell, and indicator performance is characterised. Average of 5 cells. Frame rate 20 Hz (**d**) Estimation of fluorescence-voltage relationship for the selected variant at screening. Promising NIR GEVI variants are subjected to further biophysical characterization.

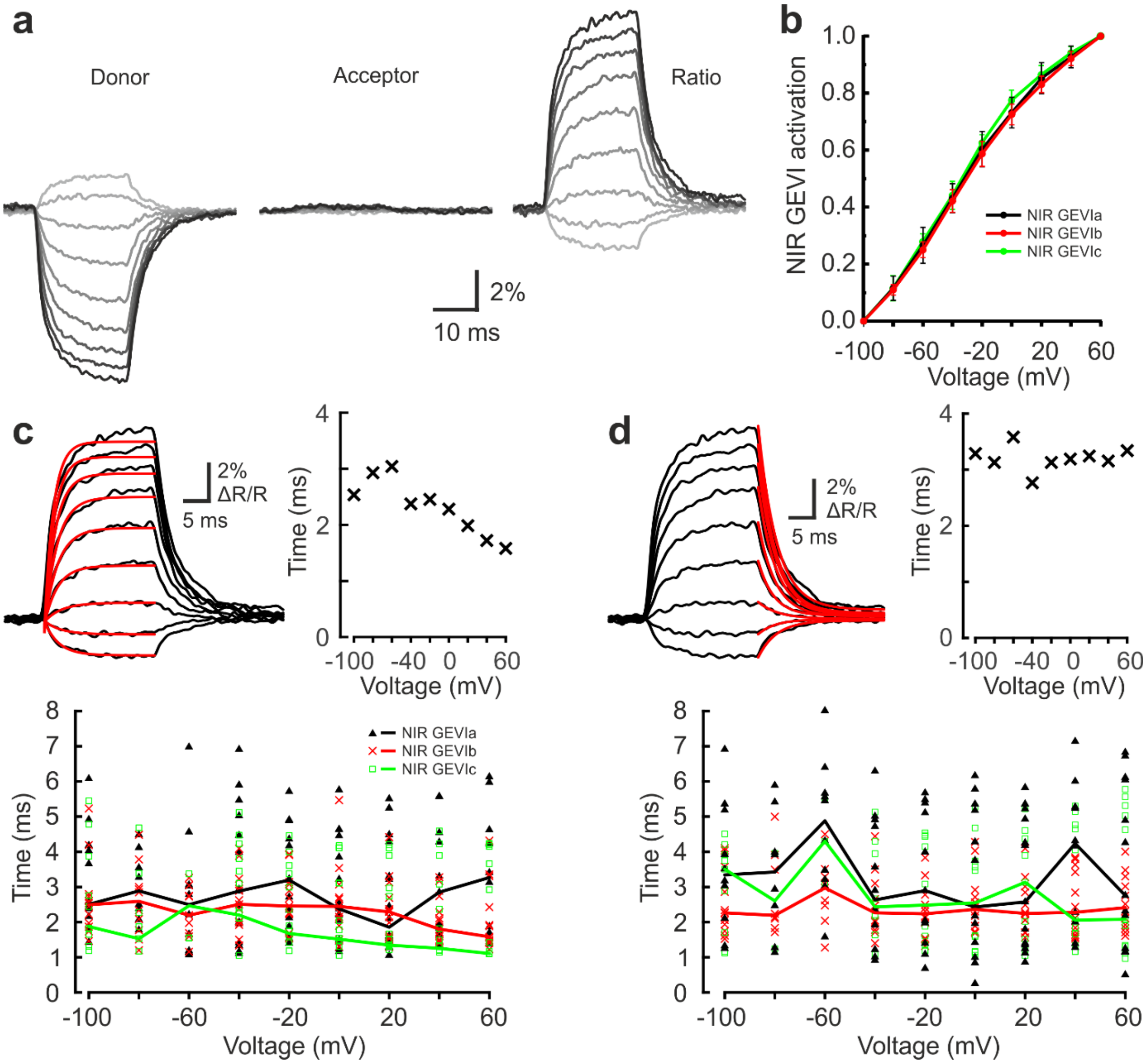

**Supplementary Figure 2.** NIR GEVI biophysical characterisation in HEK293 cells. (**a**) Fluorescence response (Donor and acceptor ΔF/F; ratio ΔR/R) to the family of voltage steps (see Online Methods) for GEVI biophysical characterisation in HEK293 cells from an example cell. Average of 10 trials. (**b**) NIR GEVIa, -b, -c) show similar activation spectra (normalised fluorescence-voltage relationship). Mean ± SD, N = 16 cells for NIR GEVIa; 15 cells for NIR GEVIb; 11 cells for NIR GEVIc. (**c**) **Upper left:** Fits of nirButterfly ON-response time constants. **Upper right**: fitted $\tau_{ON}$ for a family of voltage steps from a holding potential of -70 mV to step potentials of -100 mV to + 60 mV. **Lower**: Summary of fitted $\tau_{ON}$ for NIR GEVIa, -b, -c. Solid line indicates the median. (**d**) Same as (c) for OFF-response time constants.

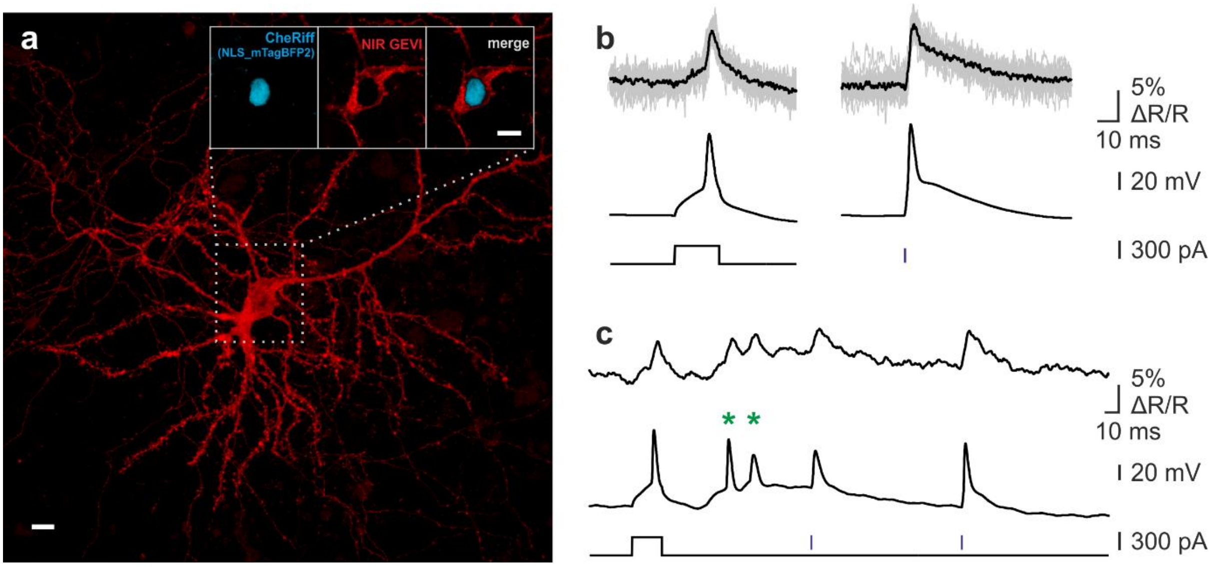

**Supplementary Figure 3.** Neurons co-expressing first generation NIR GEVIs and the cation channel opsin CheRiff also enabled all-optical electrophysiology. (**a**) Confocal image of the cultured neuron co-transfected with a NIR GEVI plasmid (NIR GEVIa) and a CheRiff-IRES-NLS-mTagBFP2, after 2% PFA fixation post-imaging (Data shown in **b** and **c**). Main image shows red channel (projection across a 9.4 um stack) and insets single red and blue channel confocal planes. Scale bar = 10 μm. (**b**) Microelectrode current injection-evoked (left) and CheRiff photocurrent-triggered (right) action potentials. Upper traces show optical signals (black: average of 19 trials; grey: single trials), middle traces show the microelectrode recording, lower traces show microelectrode (black) and blue light (blue) pulses. (**c**) Single sweep (with 50-points Savitzky-Golay filter) including spontaneous action potentials (green asterisks).

**Supplementary Table.** Main characteristics of current red-shifted GEVIs*

| GEVI name | Excitation wavelength, nm | Emission wavelength, nm | Quantum yield, % | Molecular brightness** | GEVI type | ΔF/F or ΔR/R, % | | Reference |
|---|---|---|---|---|---|---|---|---|
| | | | | | | per 100 mV | per AP | |
| FlicR1 | 568 | 592 | 49 | ~1740 | intensiometric | 9 | ~3 | [1] |
| Arch(D95N) | 585 | 687 | 0.04 | 1*** | intensiometric | ~40-60 | unknown | [2], [3] |
| QuasAr1 | 590 | 715 | 0.8 | ~20 | intensiometric | 40 | ~20 | [3] |
| QuasAr2 | 590 | 715 | 0.4 | 10 | intensiometric | 90 | ~40-50 | [3] |
| Archer1 | >650 | >664 | unknown | 10 | intensiometric | 85 | 40 | [4] |
| Archer2 | >650 | >664 | unknown | unknown | intensiometric | 60 | unknown | [4] |
| Archon1 | >637 | >664 | unknown | 28 | intensiometric | ~43 | ~30 | [5] |
| Archon2 | >637 | >664 | unknown | 80 | intensiometric | 19 | ~18 | [5] |
| nirButterfly | 640 | 670 for donor, 720 for acceptor | 14 | ~580 | ratiometric | 16 | ~4-8 | this paper |

* Red-shifted GEVIs with fluorescence emission above 550 nm are listed. **Based on the assumptions that extinction coefficient of archaeorhodopsin-based indicators is equal to that of *all-trans* retinal (52,700 $M^{-1}cm^{-1}$), and quantum yield and extinction coefficient of FlicR1 are similar to that of mApple (49% and 75,000 $M^{-1}cm^{-1}$). ***Molecular brightness (a product of excitction coefficient and quantum yield) of Arch(D95N) is assumed 1.

1. Abdelfattah, A.S., et al., *A Bright and Fast Red Fluorescent Protein Voltage Indicator That Reports Neuronal Activity in Organotypic Brain Slices.* J Neurosci, 2016. **36**(8): p. 2458-72.
2. Kralj, J.M., et al., *Optical recording of action potentials in mammalian neurons using a microbial rhodopsin.* Nat Methods, 2011. **9**(1): p. 90-5.
3. Hochbaum, D.R., et al., *All-optical electrophysiology in mammalian neurons using engineered microbial rhodopsins.* Nat Methods, 2014. **11**(8): p. 825-33.
4. Flytzanis, N.C., et al., *Archaerhodopsin variants with enhanced voltage-sensitive fluorescence in mammalian and Caenorhabditis elegans neurons.* Nat Commun, 2014. **5**: p. 4894.
5. Piatkevich, K.D., et al., *A robotic multidimensional directed evolution approach applied to fluorescent voltage reporters.* Nat Chem Biol, 2018.
